# Differential neural activity patterns mediate learning across contexts in a social cichlid fish

**DOI:** 10.1101/2021.04.04.438393

**Authors:** Mariana Rodriguez-Santiago, Alex Jordan, Hans A. Hofmann

**Affiliations:** Institute for Neuroscience, The University of Texas at Austin, Austin, Texas, USA; Department of Integrative Biology, The University of Texas at Austin, Austin, Texas, USA; Institute for Cell and Molecular Biology, The University of Texas at Austin, Austin, Texas, USA; Max Planck Institute of Animal Behavior, Konstanz, Germany

**Keywords:** animal behavior, cichlid fish, neural activity, social learning

## Abstract

Learning and decision-making are greatly influenced by the social context surrounding individuals. When navigating a complex social world, individuals must quickly ascertain where to gain important resources and which group members are useful sources of such information. Such dynamic behavioral processes require neural mechanisms that are flexible across contexts. Here we examined how the social context influences the learning response during a visual cue discrimination task and the neural activity patterns that underlie acquisition of this novel information. Using the cichlid fish, *Astatotilapia burtoni*, we show that learning of the task is faster in social groups than in a non-social context. We quantified the expression of Fos, an immediate-early gene, across candidate brain regions known to play a role in social behavior and learning, such as the putative teleost homologues of the mammalian hippocampus, basolateral amygdala, and medial amygdala/BNST complex. We found that neural activity patterns differ between social and non-social contexts. Our results suggest that while the same brain regions may be involved in the learning of a discrimination task independent of social context, activity in each region encodes specific aspects of the task based on context.

## Introduction

For group-living species, social interactions provide a key source of information that can greatly impact the fitness and well-being of individual group members. It is commonly assumed that learning from others, or *social learning*, is inherently adaptive as it allows individuals to avoid costs associated with learning by themselves, or *non-socially* [1]. The benefits of social learning allow individuals to gain information from conspecifics, such as to which foods to eat, which routes to take to feeding locations, and how to escape from predators [2]. These wide-ranging behaviors have been studied across species, such as in instances of socially transmitted food preferences [3, 4], social learning of certain skills [5, 6, 7], mate preference learning [reviewed in 8], predator avoidance [9], and fear transmission [10, reviewed in 11]. The behavioral mechanisms that underlie these behaviors are diverse, ranging from stimulus enhancement (when another individual draws the observer’s attention to a particular stimulus or object) to observational learning [12, 13, 14], allowing animals to acquire new information important for their survival and which can incidentally be transmitted to conspecifics [15, 16, 17, 18]. While a lot is known about the neural basis of learning in non-social contexts [reviewed in 19], few studies have examined whether and how these mechanisms might operate in the context of social learning.

Studies in rodents and songbirds have expanded our understanding of the neurobiological mechanisms that mediate social learning, such as the brain regions that are important for acquisition and maintenance of socially-transmitted food preferences in rats [reviewed in 20]. Subregions of the hippocampus (specifically, the subiculum and dentate gyrus) have been shown to be critical for the retention of socially acquired food preferences [21-23]. Social learning of fear has been found to be also modulated in part by the lateral nucleus of the amygdala in both rhesus monkeys and humans [11, 24].

At the molecular level, social learning requires neural activity-dependent changes in gene expression, much like long-lasting alterations in the strength of synaptic connectivity important for associative learning [25, 26]. Activation of immediate early genes (IEGs) is a critical mediator in this process [27, 28]. Previous studies in rodents have shown that IEGs such as *cfos* are expressed following acquisition and consolidation of associative learning [29 – 32]. In addition, rats trained on a test of social transmission of food preference show greater *cfos* expression in subregions of the hippocampus in a time-dependent manner [29, 30]. The medial amygdala plays a key role in mouse social cognition, as oxytocin receptors in this region are essential for recognizing familiar conspecifics [33]. In songbirds, differential Fos expression has been shown to underlie different aspects of song learning and production [34, 35]. There is also evidence in songbirds that differential neural activity underlies different phases of sexual imprinting, a type of social learning by which a juvenile learns specific characteristics of a parent or other familiar individual [36]. Taken together, these findings suggest that across species associative learning in social contexts is driven by differential neural activity patterns across multiple brain regions.

Here, we investigate the neural activity patterns that differentiate social and non-social learning in a model system that readily forms naturalistic social groups in the laboratory. The African cichlid fish, *Astatotilapia burtoni*, is a model system in social neuroscience because of its remarkable phenotypic plasticity and sophisticated social cognition [37, 38]. Dominant males of this species are territorial and aggressive, while subordinates typically do not hold territories and are overall less aggressive [38, 39]. In a recent study, we found that although dominant males of this species had strong influence over the movement of their social groups under normal conditions, they were less influential in a more complex learning task [40]. This effect was primarily driven by the socially aversive behavior of dominant males, which, although central in interaction networks, occupied peripheral positions in spatial networks. IEG expression in response to different types of social information has also been shown in this species [41 – 44], suggesting that differences in learning in social or non-social contexts may induce differential patterns of neural activity.

We examined IEG expression in different brain regions of *A. burtoni* males and females during learning in social groups or without a conspecific informant. We first compared the learning response rates in a social and non-social context as measured by the latency to acquire a cue association. We hypothesized that social facilitation mechanisms would allow groups to learn the task faster than individuals in the non-social context. To understand how the brain acquires a cued association across social contexts, we then quantified expression of Fos, an IEG, across the putative teleost homologues of the mammalian hippocampus, basolateral amygdala, and medial amygdala/bed nucleus of the stria terminalis (BNST) complex, which are key nodes of the Social Decision-Making Network (SDMN) [45, 46]. We predicted that neural activity during learning in a social context would be highest in brain regions important for mediating social behavior in this species, such as the supracommissural part of the ventral pallium (Vs, the putative homologue of the mammalian medial amygdala/BNST complex) and the medial part of the dorsal telencephalon (Dm, the putative homologue of the basolateral amygdala); as well as those important for associative learning, such as specific sub-regions of the lateral part of the dorsal telencephalon (Dl, the putative homologue of the hippocampus). In addition, we expected neural activity in Dl to increase in both contexts once learning occurs. Finally, we predicted that neural activity in regions important for social behavior would be relatively low in the non-social context. Our results reflect differences in how new information is acquired in different social contexts.

## 2. Methods

### Animals

*Astatotilapia burtoni* descended from a wild caught stock population were kept in stable naturalistic communities of eight males and eight females, as described previously [46] until being transferred to experimental aquaria. Brooding females were stripped of fry immediately prior to being placed in experimental aquaria. All work was done in compliance with the Institutional Animal care and Use Committee at The University of Texas at Austin. All relevant code and analyses are available online at https://github.com/neuromari/neuro_social_learning.

### Visual cue discrimination task

Our protocol broadly followed that of Rodriguez-Santiago, Nührenberg et al. (2020). A detailed description of the task setup, task training in a social and non-social context, as well as the response criterion we used to consider a task to have been completed successfully is provided in the Supplemental Materials. Because *A. burtoni* communities form social dominance hierarchies, we accounted for social hierarchy dynamics and group behavior in the social context, as described in Supplemental Materials.

### Sample processing and immunohistochemistry for examining neural activity

To examine neural activity patterns across learning trials, three individual samples were collected from each community. In groups with a dominant male informant, the second largest male, subordinate male, and a female were collected. In groups with a subordinate male informant, the dominant male, and third largest male, and a female were collected. For all non-socially trained individuals, males and females were euthanized after trials 6, 14, or 22. A detailed description of the immunohistochemical procedures and the quantification of Fos-positive cells is provided in the Supplemental Materials.

### Statistical analysis

All statistical analyses were conducted using R Studio (version 1.0.143) and the ‘survival’ package [47]. We analyzed the learning response using a survival analysis. We used the nonparametric log-rank test because the proportional hazard assumption was not met, given that it does not support multiple response variables, such as social context, informant status, and individual sex. Thus, we used a series of log-rank tests to examine the overall effect of social context and pairwise differences between informant status in the social context and sex in the non-social context. In a separate analysis, we examined differences between informant status effects in a social context, as well as sex differences in response rate in the non-social context using repeated measures analysis of variance (ANOVA).

We used Principal Components Analysis (PCA) to identify how neural activity patterns across brain regions clustered based on social context conditions and individual-level traits. Independent ANOVA tests were used to compare PC scores between social condition (social v non-social), trial, and learning response. To account for repeated measures of the same fish across treatments, generalized linear mixed models (GLMM) were used for Fos expression analyses, which are a proxy for neural activity. To examine how learning context, trial, and individual-level traits influence learning and neural activity patterns, we used the R package glmmTMB which ranks models based on Akaike Information Criterion scores, corrected for sample size (AICc) [48], and allows for usage of the beta family, which is appropriate for modeling proportional data. We first performed an overall GLMM that included both social and non-social learning conditions, and also did a separate model on the social and non-social conditions. In the overall model, we included learning condition, trial, sex, whether the learning criterion was met, and group as dependent variables and brain region (Dl-g, Dl-v, Dm-1, Dm-3, and Vs) as the independent variables. In the social condition model, the dependent variables were trial, sex, observer status, informant status, whether the learning criterion was met, and group. In the non-social condition model, the dependent variables were trial, sex, and learning as the dependent variables. Model results and tables can be found in the Supplemental Materials.

## 3. Results

### Social facilitation results in faster response rates compared to a non-social context

We first asked whether the cumulative response rate differed between the social and non-social contexts and found that the cumulative probability of consecutive group responses during the cue discrimination task is significantly greater than the response rate of individuals in an non-social context (log-rank test: *X*^*2*^ = 8.1, *P* = 0.004; Figure 1a). However, the number of trials it took to reach the response criterion did not differ between the social and non-social contexts (Wilcox test: W = 41, *p* = 0.426; Figure 1b). To our surprise, the social status of the informant – dominant vs. subordinate – did not have any effect on learning rate (log-rank test: *X*^*2*^ = 0.005, *P* = 0.94), contrary to our previous study [40].

**Figure 1.**
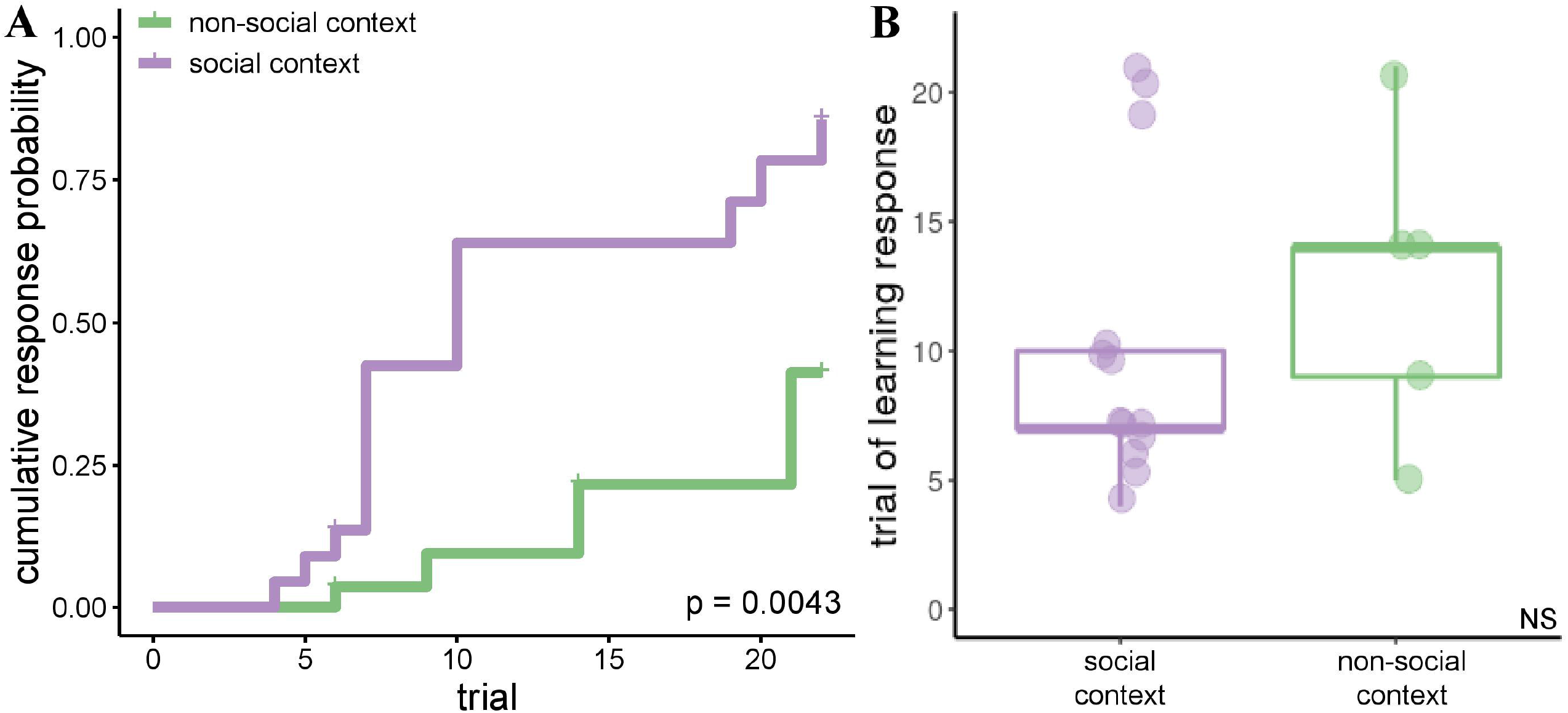
Learning rate is faster in a social context. A) Comparison of the cumulative response probability to a visual cue discrimination task between a social and a non-social context shows that groups have a higher response probability than individuals in a non-social context (p = 0.004). B) Although the rate of response probability is significantly different between social contexts, the total number of trials required to achieve this response criterion is not statistically different between contexts (p = 0.426).

### Neural activity patterns depend on the social context

We used PCA to determine which aspects of the social context and individual-level traits influence neural activity patterns during a learning task, and how these contextual aspects contribute to a learning response. We first conducted a PCA that included variables in both social conditions: the trial at which individuals were taken (trial), the context condition (social v non-social), and whether the response criterion was met (yes or no). We found that principal component 1 (PC1) accounted for 59.6% of the total variance and differed significantly between social conditions across trials (Figure 2). There was a main effect of both social context (F_1,91_ = 385.4, *p* < .001) and trial (F_2,91_ = 7.47, *p* = 0.0009), though no significant interaction effect (F_2,91_ = 1.911, *p = 0*.*154*; Figure 2d). However, there was a significant interaction between trial and learning response (learning response: F_1,91_ = 22.37, *p* < .001; trial: F_2,91_ = 3.396, *p* = .04; response x trial: F_2,91_ = 8.3, *p* < .001; Figure 2e), and strong main effects of learning response and context (learning response: F_1,91_ = 71.038, *p* < .001; context: F_1,91_ = 269.57, *p* < .001; Figure 2f). There was a strong main effect of trial and learning response on PC2, as well as interactions between trial and context, and learning response and context (Figure 2g: context: F_1,91_ = 1.086, *p* = .3; trial: F_2,91_ = 76.24, *p* < .001; interaction: F_2,91_ = 19.45, *p* < .001; Figure 2h: learning response: F_1,91_ = 59.61, *p* < .001; trial: F_2,91_ = 26.16, *p* < .001; trial x learning response: F_2,91_ = .064, *p* = .938; Figure 2i: learning response: F_1,91_ = 47.93, *p* < .001; context: F_1,91_ = 13.493, *p* = .004). Given the striking differences in neural activity patterns between the social and non-social contexts in both the comparisons of estimated Fos+ cells across brain regions and the PCA, we conducted separate PCAs on the social (Supplemental Figure 5) and non-social contexts. The results of these analysis can be found in the Supplemental Materials and Figures.

**Figure 2.**
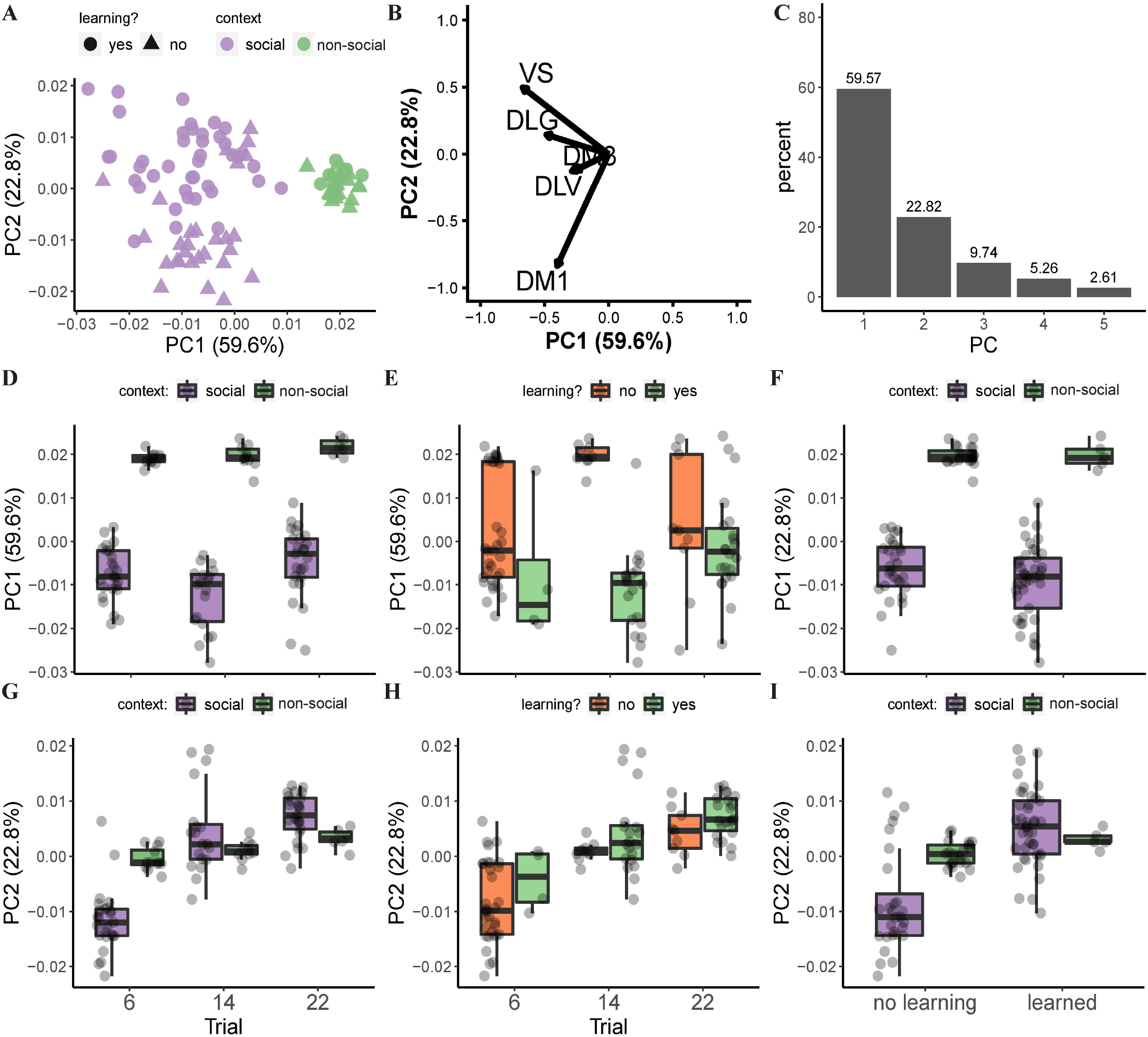
Principal component analysis (PCA) of neural activity shows differential expression pattern with learning context. A) Scatter plot of all Fos expression data separates out by social condition (social, non-social) across PC1. B) Vector plot showing the PCA variables that load on PC1 and PC2. C) Plot showing the percent of the variance explained by each PC. PC1 (D-F) and PC2 (G-I) loadings plotted across trials based on social condition (D and G) and whether learning response was reached (E and H). Boxplot showing that PC1 loadings (F) differentiates data by social condition but not by learning response while PC2 loadings (I) differentiate do not differentiate context across learning response.

### Neural activity patterns during acquisition of learning differ across social contexts

To disentangle the factors that contribute to the stark PCA differences we see with the learning context, we examined neural activity in key nodes of the SDMN across trials and contexts. We used relative Fos expression as a marker of neural activity across Dl-g, Dm-1, and Vs brain regions involved in social behavior and association learning. We compared neural activity across social conditions (social, non-social) and learning task trial (6, 14, 22) using two-way ANOVAs (Figure 3a,c,e; Supplemental Table 3 for statistics). We found that the trial and context had significant main effects on Fos expression in the Dl-g, but there was no significant interaction. In the Dm-1, there was both a significant main effect of trial and context as well as an interaction. In the Vs, there was a main effect of trial and context. In addition, we also examined neural activity in the Dl-v and the Dm-3 subregions and found no significant effect of trial, although there was a significant effect of context (data not shown, statistics in Supplemental Table 3).

**Figure 3.**
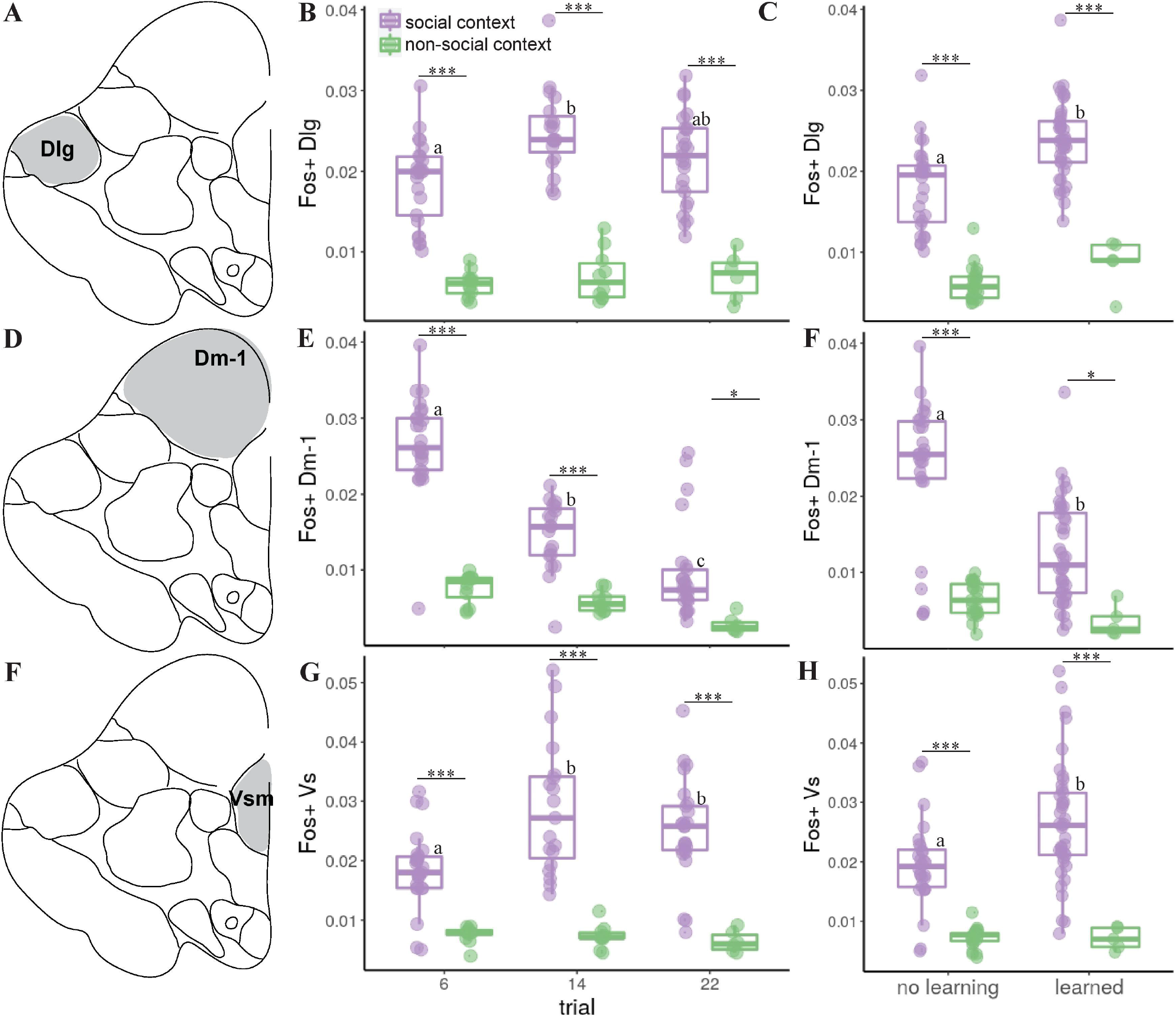
Neural activity across brain regions varies over trials and with learning. Fos expression was quantified as a marker of neural activity in the Dlg, Dm-1, and Vs regions of the forebrain (A, D, F). In the Dlg, there was a significant increase in activity from trial 6 to 14 in the social context, while there was no difference in activity across trials in the non-social context (B). Neural activity was significantly highest in learners in the social context (C). In the Dm-1, activity significantly decreased over trials (E). Activity was significantly highest in the Dm-1 in the social context when learning had not occurred (F). In the Vs, activity significantly increased after trial 6 in the social context (G) and was significantly higher in the social context with learning (H).

When we examined whether Fos expression changed with learning, we found a significant difference between context treatments (Figure 3b,d,f; see Supplemental Table 4 for all statistics). Across all brain regions, there was a main effect of context and learning response. There was an interaction between learning and context in the Dm-1 only. There was no difference in Fos expression in the non-social context based on whether individuals learned the task, while in the social context Fos expression was highest in observers that learned the task in the Dl-g (*p <* 0.001; Figure 3b) as well as in the Vs (*p* = 0.001; Figure 3f). Despite the large differences between Fos expression across social contexts in the five brain regions measured over trials, when we looked closer at factors that impact this difference within the social context we found no significant differences in expression based on the social rank of observers or based on the informant status (data not shown).

## 4. Discussion

In the present study, we found large differences between a social and non-social context in behavioral and neural activity during an associative learning task. Specifically, we discovered a significant difference in learning rate between contexts, such that social groups had a higher cumulative probability of reaching the response criterion sooner than individuals in a non-social context. This striking behavioral difference is reflected in the neural activity pattern differences between contexts, with specific brain regions encoding different aspects of our learning paradigm, suggesting that the acquisition of a learning response to a cue association is mediated by different brain regions depending on the social context.

### Observational learning and stimulus enhancement accelerate associative learning of a visual cue discrimination task

By examining the cumulative learning rate probability of acquiring a cue association response across two experimental contexts, we found that social groups had a significantly higher cumulative probability of learning than individuals in a non-social context. This is not necessarily surprising given the prevalence of social learning strategies across species and the notion that social learning is more adaptively beneficial as it confers fewer costs and allows individuals to gain new information more quickly [49]. In addition, information diffusion is typically accelerated in social groups [20].

There are at least two mechanisms by which learning might have occurred in our social paradigm, social facilitation (when the presence of an informant affects the observer’s behavior) and stimulus enhancement (where the observer’s behavior changes after watching an informant interact with a stimulus). To demonstrate that the group response is due solely to the presence of an informant (i.e., social facilitation), it would be necessary to test individual group members by themselves following acquisition. While we did not examine this retention by observers in the present study, it should be noted that the informants themselves were trained in naïve groups and then transplanted to new groups, where they were the only informed individual. Importantly, all informants displayed a correct response to the cue within one or two trials in their new communities, suggesting that the observers in our study in fact acquired the association and were not just copying other group members’ behavior. It seems thus likely that individuals in social groups learned by means of observation or stimulus enhancement, which ultimately led them to respond faster than lone individuals. However, it cannot be ignored that *A. burtoni* is a highly social species, and although individuals in the non-social context had blind cave fish as a social buffer, their slow learning rate could be due to stress factors from being apart from conspecifics.

### Hippocampal sub-regions differentially mediate learning in social and non-social contexts

When we examined the neural activity across brain regions in different trials of the learning task, we found significant differences in neural activity (measured as number Fos-immunoreactive cells) between the social and non-social contexts as well as depending on whether the task had been learned or not. In Dl (the lateral part of the dorsal telencephalon and putative homologue of the mammalian hippocampus), we found a significant increase in Fos expression (or ‘activity’) from trial 6 to trial 14 in the social context in the Dl-g sub-region, which was also significantly higher in groups that learned the task. In the Dl-v sub-region, there was a significant main effect of social context across trials, and a significant decrease in activity between trials 14 and 22. Activity in the Dl-v was not correlated with learning. The Dl-g and Dl-v are subdivisions of the dorsal pallium, a region implicated in the learning of spatial and temporal relationships in teleosts [50, 51]. Previous work has also shown that the major pathways within the dorsal pallium are highly recursive and have complex reciprocal connections with subpallial regions [52]. Based on tract-tracing neuroanatomical data, as well as lesions studies that implicate the Dl and other dorsal pallial regions in learning and memory tasks, Elliott et al. (2016) suggested that the dorsal pallial circuitry (which includes the Dl subregions) can implement the same pattern separation and completion computations ascribed to the mammalian hippocampal dentate gyrus and CA3 fields. Taken together, these results suggest a differential role for these Dl subregions in the acquisition of this association learning task.

### The basolateral amygdala likely encodes social group formation, not learning of the association task

We found a significant difference in neural activity across social contexts in subregions of Dm (medial part of the dorsal telencephalon and putative homologue of the mammalian basolateral amygdala). More specifically, we found a significant decrease in activity across trials in the social context in Dm-1, and a significant decrease from trial 14 to 22. Activity in Dm-1 was not associated with group learning, and there was no difference across learning response in the Dm-3 (not shown). These findings are consistent with previous studies in goldfish that have shown that Dm lesions disrupt trace and delay avoidance conditioning [51, 53], as well as fear and heart-rate classical conditioning [22], while such lesions have no effect on spatial memory and cue learning [54, 55]. The effects of these lesions in fish are similar to lesions of the amygdala in mammals [56 – 59] and in part based on this evidence the teleost medial pallium (which includes the Dm) has been proposed as homologous to the pallial amygdala of mammals [51]. In *A. burtoni* males, Dm activity is correlated with the level of engagement in joint territory defense, although the sign of the correlation depends on an individual’s role in this cooperative behavior [44]. In the present study, we found that activity in the Dm complex was significantly higher in trial 6 compared to 14 and 22. Given that few groups had learned the task prior or by trial 6, it is not surprising that Dm activity was also higher in groups that had not yet successfully learned the task. Interestingly, individuals from groups that did reach the learning criterion by trial 6 or sooner showed lower Dm activity, which further indicates that Dm is not involved in learning the cue association task. Instead, this result suggests that the Dm regions, and the Dm-1 in particular, may play a role in some aspect of social group formation rather than being involved in the acquisition of the cue association task, providing further support for a role of this brain region in affective processing.

### The extended medial amygdala encodes social context

In Vs (the supracommissural part of the ventral pallium and putative homologue of the mammalian medial amygdala/BNST) we found a significant main effect of social context. Also, Vs activity increased in social groups in trials 14 and 22, possibly as a consequence of more groups successfully learning the task at these later trials. Homology of this brain region has historically been difficult to characterize due to the eversion, rather than invagination, of the neural tube during teleost development [60 – 63]. However, developmental studies have found similar genetic markers, namely *Dlx2, Lhx7, Nkx2*.*1b*, between the Vs and the extended amygdala [64]. Stimulation of the Vs has been shown to increase aggression in male bluegill fish [65]. In our species, *A. burtoni*, this region is under social and reproductive modulation [42] and shows varying levels of sex steroid receptor expression in males when given the opportunity to ascend or descend in status. Taken together, this suggests that Vs plays a predominant role in mediating social information, which is why we see large differences in neural activity here between the social and non-social learning contexts.

### Disentangling the effects of group formation and learning on neural activity patterns

While we see evidence for differential neural activity across multiple brain regions during the acquisition of an association in both social and non-social contexts, we are unable to fully separate the effects of group formation time from the effects of learning. Even though there are significant differences in neural activity in specific brain regions (Dl, Dm) based on whether groups demonstrated learning, it remains unclear how group formation impacts learning. In other words, there could be a dampening of response in early trials due to social instability simply because the groups did not have time to acclimate prior to the start of the trials. In the non-social context, we observed a general dampening of neural activity specifically in early trials that coincided with lower behavioral activity levels. Disentangling the effects of social stability formation from the increased probability of learning after repeated trials in both social and non-social contexts will require subsequent rigorous behavioral examination with automated tracking.

### What Fos expression tells us about the observed neural activity patterns

An important aspect of examining IEG induction as a measure of neural activity is that we examined this expression 1 hour after the last learning trial the animals underwent – whether it was trial 6, 14, or 22. Expression of IEGs such as Fos is widely used as a measure of neural activity [66,67] as most IEGs encode transcription factors or DNA-binding proteins that coordinate the cellular response to a stimulus [28]. By examining Fos protein expression within 60-90 minutes following the last stimulus exposure, we aimed to capture the brain regions that were active, and presumably important, for the animal’s behavioral response. Animals did not perform these behaviors in isolation, and it is possible that both in the social and non-social contexts their neural activity reflects a response to the environment rather than the stimulus cue itself. For example, there could have been a salient social signal occurring in the aquarium at the same time as the cue (such as high territorial aggression by a dominant male). However, given the high Fos expression in the Dm-1 in trial 6 compared to later trials in both the social and non-social contexts, the observed IEG pattern in this region is likely reflective of the animal’s response to other salient cues in the (social) environment besides the stimulus cue. In addition, we found no correlation between neural activity and informant aggression (data not shown), although the aggressive behaviors of other observer males could have had an effect on the neural activity patterns seen in the social context.

### Group learning and neural activity patterns are independent of social status

Communities of *A. burtoni* naturally form rank hierarchies with some males establishing social dominance by aggressively defending territories for mating with females, while the majority of males are socially subordinate and reproductively suppressed [37, 68]. We have previously shown for this species that the social status of an informant can have a strong effect on how fast a group learns the visual cue discrimination task we used in the present study. Specifically, even though socially dominant males strongly influence their social groups through aggressive displays and space use, they are significantly less effective in generating group consensus during the association task than subordinate males [40]. In contrast, we did not find a significant effect of informant status on group learning in the present study. This may not be surprising given that the present study was not designed to examine the effects of social status on group learning, and thus lacks the statistical power to robustly detect such an effect. It should also be noted that in the Rodriguez-Santiago, Nührenberg et al. (2020) study, dominant males were considerably larger than subordinate males, while in the present study the size difference was much smaller. Previous work has shown that small size differences result in lower stability of the social hierarchy in this species [69]. Although we did not quantify group stability here, the behavioral traits that determine whether an individual is an effective informant – aggression and space use – are highly context-specific and might explain the absence of a social status effect. These factors may also explain why we did not find differences in neural activity patterns between dominant and subordinate observers when learning the task. One interesting observation of relevance here comes from social fear learning in rats, where subordinate animals display increased fear responses after interacting with a dominant informant, which is also reflected in distinct neural activity patterns [70].

## 5. Conclusion

We used the highly social African cichlid fish *A. burtoni* to demonstrate that social learning is associated with increased neural activity (as measured by the expression of Fos, an IEG) when compared to non-social learning across key brain regions important for learning and social behavior. These brain regions are important for modulating learning (hippocampus), emotional learning and fear avoidance (basolateral amygdala), and social behavior (medial amygdala/BNST), and are part of a greater Social Decision-Making Network that is important for mediating various aspects of social behavior [45, 46]. In addition, we found that activity in these regions was not modulated by the sex or social status of individuals, nor was it impacted by the status of informants in social groups. Thus, while these regions are important for different aspects of social learning [45], they do not appear to be modulated by group dynamics or individual-level traits in a social learning context. While future studies are needed to fully understand the mechanisms that drive social learning contexts (e.g. neuroendocrine or dopaminergic pathways), our results in *A. burtoni* highlight that there are neural activity pattern differences in how individuals acquire information in different social contexts.

## 6. Acknowledgements

We thank members of the Hofmann lab for many fruitful discussions. In particular, we thank Caitlin Friesen and Isaac Miller-Crews for comments on earlier drafts of this manuscript, and Nupur Shambharkar for performing the Fos cell counts. In addition, we thank Julie Butler and Morgan Gustison for detailed comments on earlier drafts of this manuscript. This work was supported by a UT Austin Graduate School Bruton and Summer Fellowships, and a Department of Integrative Biology Doctoral Dissertation Improvement Grant (MRS); the National Science Foundation Bio/computational Evolution in Action Consortium (BEACON) Center for the Study of Evolution in Action (H.A.H. and A.J.), Dr. Dan Bolnick and the Howard Hughes Medical Institute (A.J.); and NSF grant IOS1354942 (HAH).

## 7. Author Contributions

MRS, AJ, and HAH designed experiments, MRS performed experiments and statistical analysis, MRS and HAH wrote the manuscript, MRS, AJ, and HAH revised manuscript.

