## Supplemental Materials and Data for "Differential neural activity patterns mediate learning across contexts in a social cichlid fish"

*Visual cue discrimination task*

In a protocol broadly following that of Rodriguez-Santiago et al. (2020), all animals were housed and tested in 200 L aquaria (108 x 54 x 46 cm) with digital video cameras mounted above (see Supplemental Figure 1 for schematic). Cameras were scheduled to record behavior for 20 minutes prior to each trial and for 10 minutes after. Each aquarium had two automatic feeders (Eheim) attached at opposing ends. Each pair of feeders was controlled by an Arduino Uno microcontroller board attached to an LED, which presented either an orange or cyan light for three seconds every three hours. Following the offset of the LED cues, the feeder programmed to shine the orange light immediately dispensed food (cue + feeding sides were randomized across trials to ensure the animals did not develop a side preference and that there was no local enhancement). The first cue was at 8:00h every day, for a total of four cue trials per day every three hours. Food reward was dispensed after every cue regardless of behavioral response. This was the learning task used for training informants (see below), as well as the social groups with informants and the individuals in the non-social context (detailed below).

*Training informants on the discrimination task*

Twenty males and twenty females were tagged with visible implant elastomer tags (Northwest Marine Technology) for identification one week prior to the start of the experiment. Social groups of four tagged males and four tagged and non-gravid females (n = 5 groups) were initially placed together in the experimental aquaria 16 hours prior to the first learning trial. Following twenty-two trials, individuals were placed in holding aquaria and remained in the same social groups in which they were trained. These individuals are now considered *informants* for observers in the subsequent learning experiments.

Following informant training, one male (n=11 dominant male informants, n=11 subordinate male informants) was placed in a new group of all naïve fish (deemed *observers*). Groups of seven observers (three males, four females) and one informant male were trained on the visual discrimination task described above (n=11 total groups with dominant informant, n=11 total groups with subordinate informant). Every group (including the informant) consisted of four males and four females (females measured on average 41.5±0.55 mm and were within 5 mm length of other females in the tank). Groups were assembled and placed in the experimental aquaria 15 hours prior to the first trial at 8:00h the following day. Groups were trained for either 6, 14, or 22 trials (depending on their pre-determined experimental group). Three naïve study subjects (the largest (51±1.3 mm) and smallest (48.5±1.2 mm) male observers, and one female) were taken from each group after either trial 6, 14, or 22 for subsequent brain collection and histological processing. The last trial (either 6, 14, or 22) for all groups was at 11:00h.

We examined the aggressive behavior of informants the ten minutes prior to every trial by quantifying approach behavior, defined as the focal informant swimming directly towards any part of another fish’s body, within 3 body lengths. If the approached fish responded by moving away in any direction, the behavior was recorded as a displacement for the focal initiator.

*Visual cue discrimination task training in a non-social context*

In the non-social condition, naïve individual males (n=14) and individual females (n=14) were trained on the same visual discrimination task while housed with seven blind cave fish (Mexican tetra, *Astyanax mexicanus*) such that the focal fish is socially stimulated but receives no social facilitation while responding to the visual cue. All males tested were dominant as all males transitioned into the dominant phenotype immediately after being put in isolation. Individuals were placed in the experimental aquaria along with blind cave tetras the night before the first trial. All individuals were taken after either trial 6, 14, or 22 for subsequent brain collection and histology.

*Response criterion for successful task completion*

For each behavioral trial, we assessed the response of individual fish to the onset of the LED and quantified the number of fish that moved toward the correct feeder prior to the food reward being dispensed. For all fish, a correct response required the fish to move directly under the correct feeder in order to be scored. In the social context, the percent group response was calculated by dividing the number of fish that moved by the total number in the group. In the non-social context, individual response was binary – either the fish moved or did not. We established a response criterion where groups and individuals had to correctly respond to the task in two or more consecutive trials for them to be considered to have responded correctly, or *learned* the task. For the social context, this criterion was met when at least five of the seven naïve group members (71%) move to the correct cue. Note that this variable is right censored, because we stopped the experiment after either six, fourteen, or twenty-two trials.

*Learning Response Sensitivity Analysis*

We tested the sensitivity of this response criterion by examining how changes to the response parameters impacted the cumulative response rate (Supplemental Figure 2). Specifically, we used survival analysis to compare the response rates of 50-100% of the group in two, three, and four consecutive trials. We found no significant difference in the cumulative response probability (Supplemental Table 1) of the group when we compared each required group percent response after two, three, or four consecutive responses. When we examined how the cumulative response probability differs depending on the percent of group members that responded across trials, we found a significant difference (log-rank test: *X^2^* = 42.7, *P* < 0.001; Supplemental Figure 2). For the non-social context, we tested the response criterion by examining how the results are affected by changing the number of consecutive trials required to reach criterion. We found that there is a significant difference in response probability between two, three, and four consecutive trials as the response criterion (log-rank test: *X^2^* = 6.1, *P* = 0.05; Supplemental Figure 2d). Based on these validation experiments, we concluded that using a group response criterion of 70% after two or more consecutive trials was most biologically meaningful since it does not show a response floor or ceiling effect.

*Accounting for social hierarchy dynamics and group behavior in the social context*

To understand the social factors that might contribute to variation in informant influence, we examined both the aggressive behavior of informants as well as the total behavioral activity of the entire community. To determine whether informants retained their social status over time, we examined the number of displacements as a measure of their aggressive behavior). A repeated-measures analysis of variance (ANOVA) indicated that the number of displacements did not change over time (F_21,239_= 0.91, p = 0.579), demonstrating that informants maintained their social status throughout the experiment (Supplemental Figure 3a). As expected, there was a significant effect of informant status (F_1,279_ = 22.229, p < 0.001), confirming that DOM and SUB informants differed in their overall aggression. Importantly, however, the number of displacements by an informant (as a proxy of aggression) did not affect the number of trials it took the group to reach criterion (*r* =0.65, F_21,281_ = 0.646, *p* = 0.882; Supplemental Figure 3b).

We also examined the relationship between behavioral activity and learning response in both social and non-social treatments (Supplemental Figure 4). We found that overall behavioral activity (as measured by total number of line crosses in the 10 minutes prior to the trial, normalized to the number of group members) increased in both treatments over time (F_1,42_ = 17.645, *p* = <0.001), although there was no interaction between treatment (social v. non-social) and behavioral activity over trials (F_3,42_ = 2.041, *p* = 0.123; Supplemental Figure 4a). Although social groups had higher overall behavioral activity than the non-social context, this was not driven by the number of displacements by informants (F_1,301_ = 0.084, *p* = 0.77; Supplemental Figure 4b). In addition, although there was a significant increase in mean behavioral activity over trials across all social groups, there was no main effect of learning rate or interaction between learning rate and trial on behavioral activity (ME of learning rate: F_2,16_ = 0.497, *p*=0.618, learning rate x trial: F_2,16_ = 0.289, *p* = 0.753; Supplemental Figure 4c). In the non-social context treatment, there was a main effect of trial and learning rate on mean behavioral activity (trial: F_1,23_ = 15.156, *p* = 0.001, learning rate: F_2,23_ = 11.638, *p* < 0.001), but no significant interaction between learning rate and trial (F_1,23_ = 1.044, *p* = 0.317; Supplemental Figure 4d).

*Sample processing for examining neural activity and Quantification of Fos-positive cells*

To examine neural activity patterns across learning trials, three individual samples were collected from each community. In groups with a dominant male informant, the second largest male, subordinate male, and a female were collected. In groups with a subordinate male informant, the dominant male, and third largest male, and a female were collected. For all non-socially trained individuals, males and females were euthanized after trials 6, 14, or 22 (see Supplemental Table 2 for sample sizes).

Brains were rapidly dissected, fixed overnight in 4% paraformaldehyde at 4°C, then washed in 1X phosphate-buffered saline (PBS) and cryo-protected in 30% sucrose at 4°C, then embedded in O.C.T. compound (Tissue-Tek; Fisher Scientific Co., Pittsburgh, PA, USA), and stored at -80°C until further processing. Brains were sectioned on a cryostat at 30 μm and thaw-mounted onto Super-Frost Plus slides (Fisher Scientific) in four series of alternating sections that were stored at -80°C until further processing. For Brightfield detection of the IEG Fos, one series of sections was removed from -80°C, dried on a slide warmer and processed for immuno-histochemistry (IHC), as described previously (Weitekamp et al., 2017), using a rabbit anti-c-Fos primary antibody (Santa Cruz Biotechnology, Inc., Santa Cruz, CA; catalog #sc-253; 1:500 dilution).

Slides were coded such that the experimenter was blind to treatment. Cell nuclei labeled by Fos IHC were clearly marked by dark brown staining and counted using the Optical Fractionator workflow of the Stereo Investigator software package (Microbrighfield, Williston, VT, USA). We quantified Fos induction in two subregions, Dm-1 and Dm-3, of the medial part of the dorsal telencephalon (Dm, the putative teleost homolog of the mammalian basolateral amygdala), the supracommissural nucleus of the ventral pallium (Vs, the teleost homolog of the medial amygdala/bed nucleus of the stria terminalis), and the lateral subdivision (Dlv) and the granular zone (Dlg) of the lateral part of the dorsal telencephalon (Dl, the putative teleost homolog of the hippocampus) (O’Connell & Hofmann, 2011). Three sections of each brain region were quantified and averaged per individual. A region of interest was defined using a 2x objective, then positive cells were counted using a 20x objective. The counting frame and sampling grid parameters varied per brain region to account for differences in cell density and overall area (Vs: 30x30 counting frame, 100x100 sampling grid; Dm, Dlv, Dlg: 50x50 counting frame, 150x150 sampling grid). For each brain region, data are presented as the estimated population of Fos immunoreactive nuclei using number weighted section thickness divided by the area of the region.

To examine how neural activity patterns change in the social context during the acquisition of a cue association, we counted Fos expression in key brain regions of interest, namely the ventral (Dl-v) and granular (Dl-g) subregions of the lateral part of the dorsal telencephalon (Dl), the Dm-1 and Dm-3 subregions of medial part of the dorsal telencephalon (Dm), and the ventral pallium (Vs) (O’Connell & Hofmann, 2011). We first confirmed that there was not a correlation between fish standard length (in mm) and IEG positive cell numbers in any brain region across both contexts (data not shown). Next, we compared how activity in these regions changed across trials in different social conditions (non-social / individuals trained by themselves, and social / observers trained with an informant in groups). For the non-social context, we further examined how this activity across trials was influenced by the animal’s sex. For the social context, we examined the impact of informant social status (dominant or subordinate males) as well the sex and size of the observers.

*Neural activity patterns vary depending on the social context*

Given the striking differences in neural activity patterns between the social and non-social contexts in both the comparisons of estimated Fos+ cells across brain regions and the PCA, we conducted separate PCAs on the social (Supplemental Figure 5) and non-social contexts (Supplemental Figure 6). In this analysis, we found that PC1 accounts for 43.98% of the total variance in the social context, and while there was not a clear separation by informant status (Supplemental Figure 5a), Dm-1 and Vs regions loaded most strongly on PC1. There was a main effect of trial and learning response (trial: F_2,63_ = 19.43, *p* < .001; learning response: F_1,63_ = 77.28, *p* < .001; Supplemental Figure 5e) but no significant main effect of informant status on PC1 (F_1,63_ = 1.525, *p* = 0.221) (Supplemental Figure 5d). There was not a significant interaction between learning response and informant status on PC1 in the social context (F_2,63_ = 1.889, *p* = 0.160) (Supplemental Figure 5f). In addition, there was a main effect of trial on PC2 (F_2,63_ = 6.168, *p* = .003), but no effect of informant status or learning response (Supplemental Figure 5g-i). In the non-social context, we found similar striking differences in the brain regions that loaded on PC1 (53.97% of the total variance) and PC2 (31% of the total variance) (Supplemental Figure 6b). There was a main effect of learning on PC1 (F_1,22_ = 7.732, *p* = .011; Supplemental Figure 6e-f) but no main effect of sex or trial. For PC2 there was a main effect of trial (F_2,22_ = 10.22, *p* < .001; Supplemental Figure 6g-h) but not main effect of sex or learning.

*Experimental variables drive neural activity in a brain region-specific and social context-dependent manner*

We employed generalized linear mixed models and AIC_c_ scores to understand which variables influenced activity in each brain region given that the PCA results show that differential brain region activity underlies learning across contexts (Figure 2). We compared the AIC_c_ scores to pick the model that best represents the variables that influence activity in each brain region within each social context. The first models for each social condition included all variables accounted for in each context (social context included: informant status, observer status, trial, learning response; non-social context included: sex, trial, learning response). For the Dl-g, we found that the learning response in the model of best fit for both the social and non-social context had the greatest effect on activity (social: AIC = -532.3, *p* < .001; non-social: AIC = 259.1, *p* = .049). Meanwhile in the Dm-1, activity in both contexts was mostly driven by trial (social: AIC = -504.5, *p* < .001 for both trials 14 and 22; non-social: AIC =278.2, *p* = .018 for trial 14, *p* < .001 for trial 22). For the Vs, activity in the model of best fit was driven by trial 14 in the social context (AIC = 468.3, *p* = .004), while there was no effect of any predictor variable in the non-social context (for full statistical models, see Supplemental Table 5 for model comparison in just the social context, and see Table 6 for model comparison in just the non-social context). Together, these data suggest that neural activity across social conditions was predominantly driven by factors that were not part of the social context, such as whether individuals learned the task and the task trial.

**Supplemental Figures**

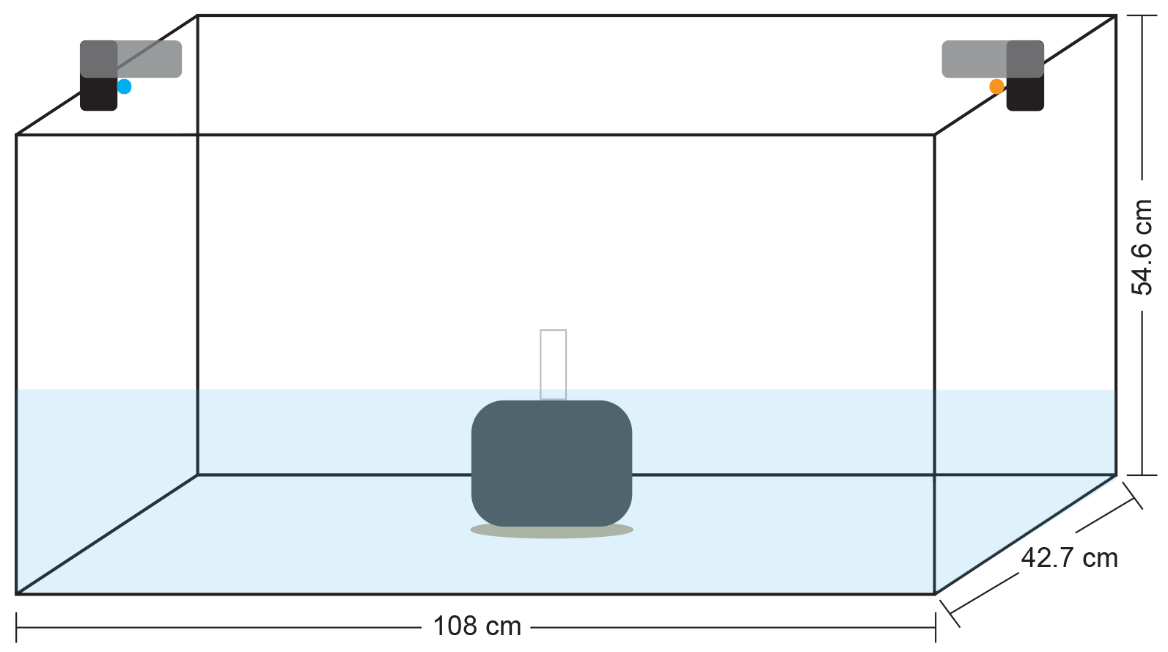

*Supplemental Figure 1*. Schematic of aquaria used for housing during the learning experiment. All animals were housed in these aquaria for either two, four, or six days depending on their experimental condition. Automated feeders were connected to Arduino Uno boards and placed at the top of the aquaria such that LED lights shined colors down into the water. All aquaria had an air filter in the center.

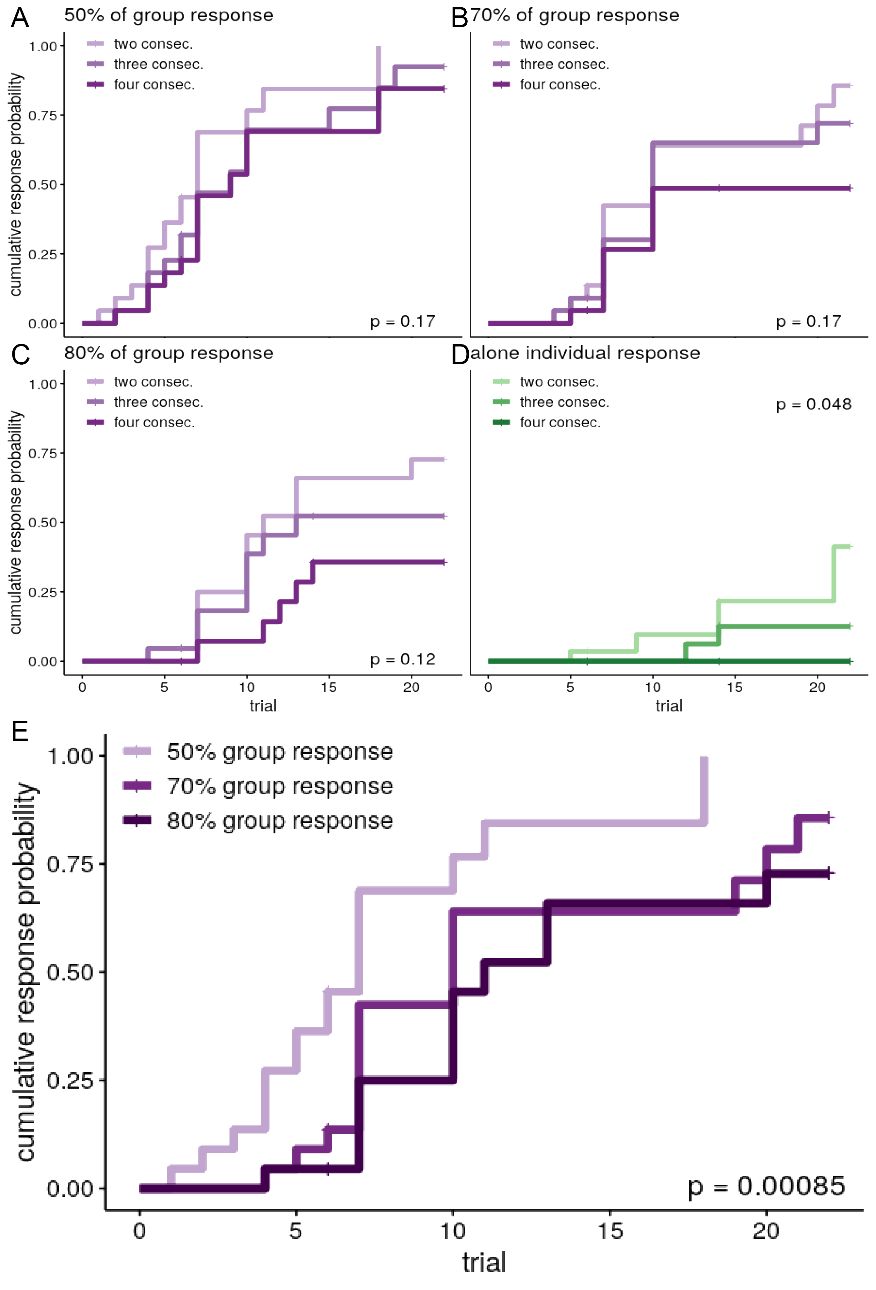

*Supplemental Figure 2*. Behavioral sensitivity of cumulative response probability across learning context. A-C) Comparison of the cumulative response probability to a visual cue discrimination task between 50, 70, and 80% of the group is not significantly different between number of consecutive trials required for response. D) There is a significant different between the cumulative response probability in the non-social context between number of consecutive trials required to meet the response criterion, though the response probability never reaches 0.5 (or 50%). E) In the social context, there is a significant difference in the response probability based on the percentage of the group that must respond in two consecutive trials (*p* < 0.001).

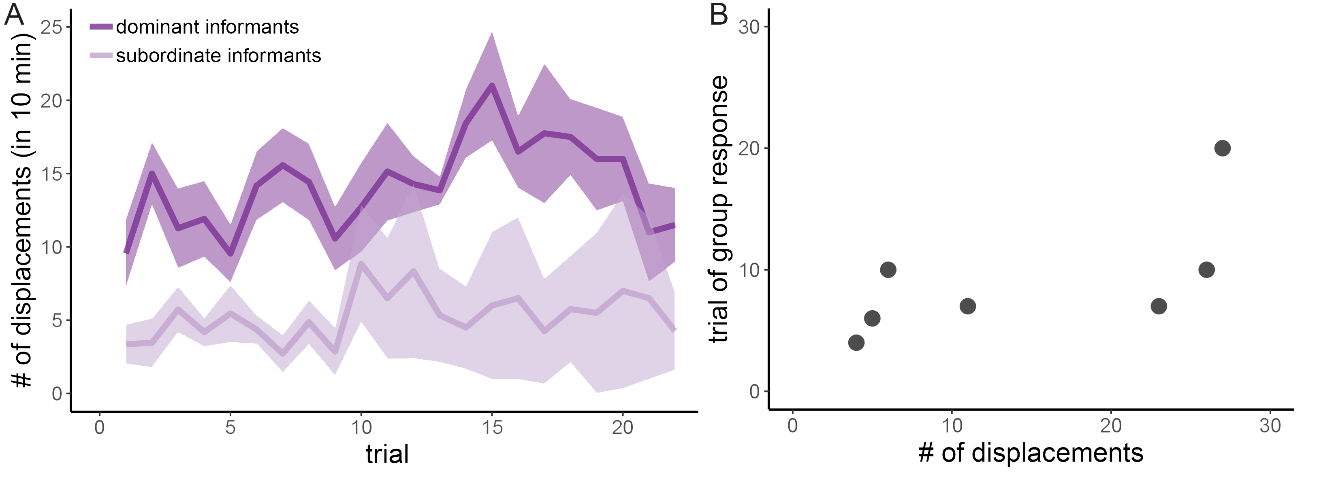

*Supplemental Figure 3*. The impact of informant aggression on social group response. A) The number of aggressive displacements by informants in the 10 minutes prior to each trial did not change over time, although there was a significant difference in overall displacements between dominant and subordinate informants. B) Although there is a positive correlation between the number of displacements by an informant at the trial a social group reached criterion, this relationship is not significant. Overall this indicates that although dominant and subordinate male informants differed significantly in their aggression, their aggressive behavior did not influence the trial at which social groups reached response criterion.

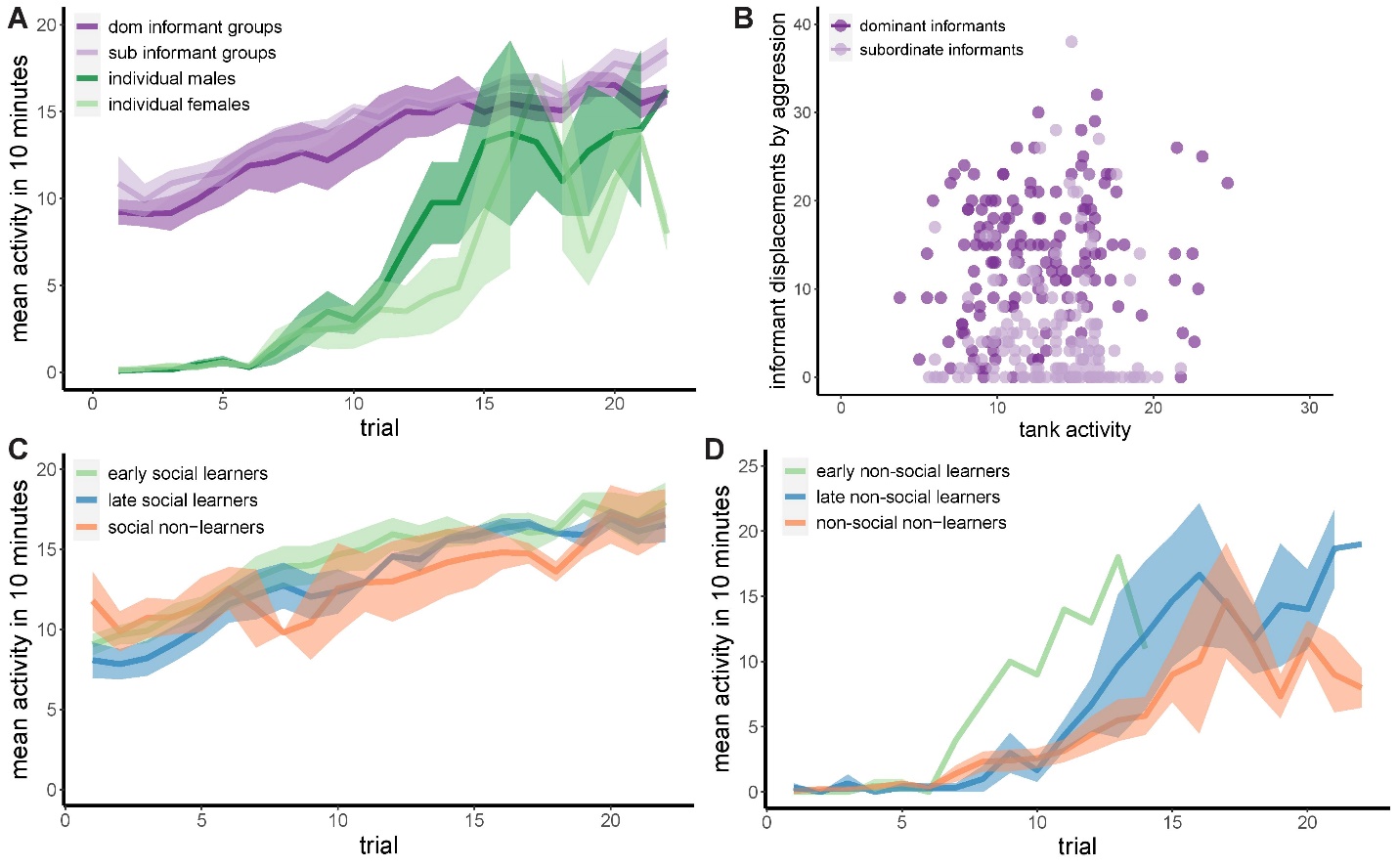

*Supplemental Figure 4*. The impact of behavioral activity on learning. A) Across all treatments, the mean behavioral activity (measured as line crosses in the 10 minutes prior to each trial) significantly increased over time and was overall higher in social groups compared to the non-social condition. B) There was no significant relationship between social group activity and informant displacements, demonstrating that higher behavioral activity in the social groups was not driven by the aggressive behavior of informants. C) There was no significant difference between learning rate on social group activity over trials. D) There was a main effect of learning rate on mean behavioral activity in the non-social condition across trials.

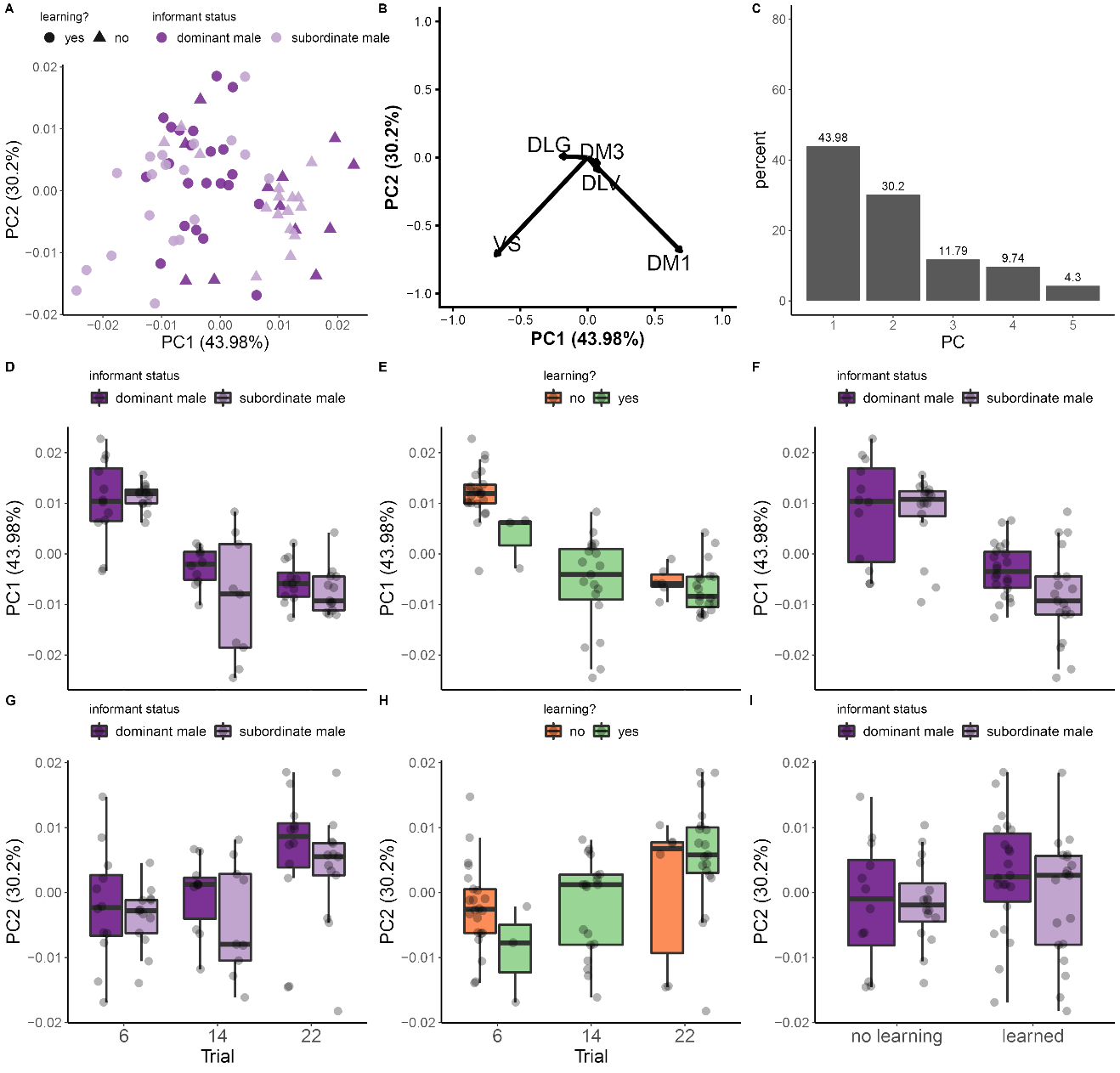

*Supplemental Figure 5*. Principal component analysis (PCA) of neural activity patterns in a social group context. A) Scatter plot of Fos expression data does not pull out differences by informant status across either PC, however PC1 separated out the Dm-1 and Vs brain regions (B). C) Plot showing the percent of the variance explained by each PC. PC1 (D-F) and PC2 (G-I) loadings plotted across trials based on informant status (D and G) and whether learning response was reached (E and H). Boxplot showing that PC1 loadings (F) differentiates data by learning response but not by informant status while PC2 loadings (I) differentiate do not differentiate context across learning response.

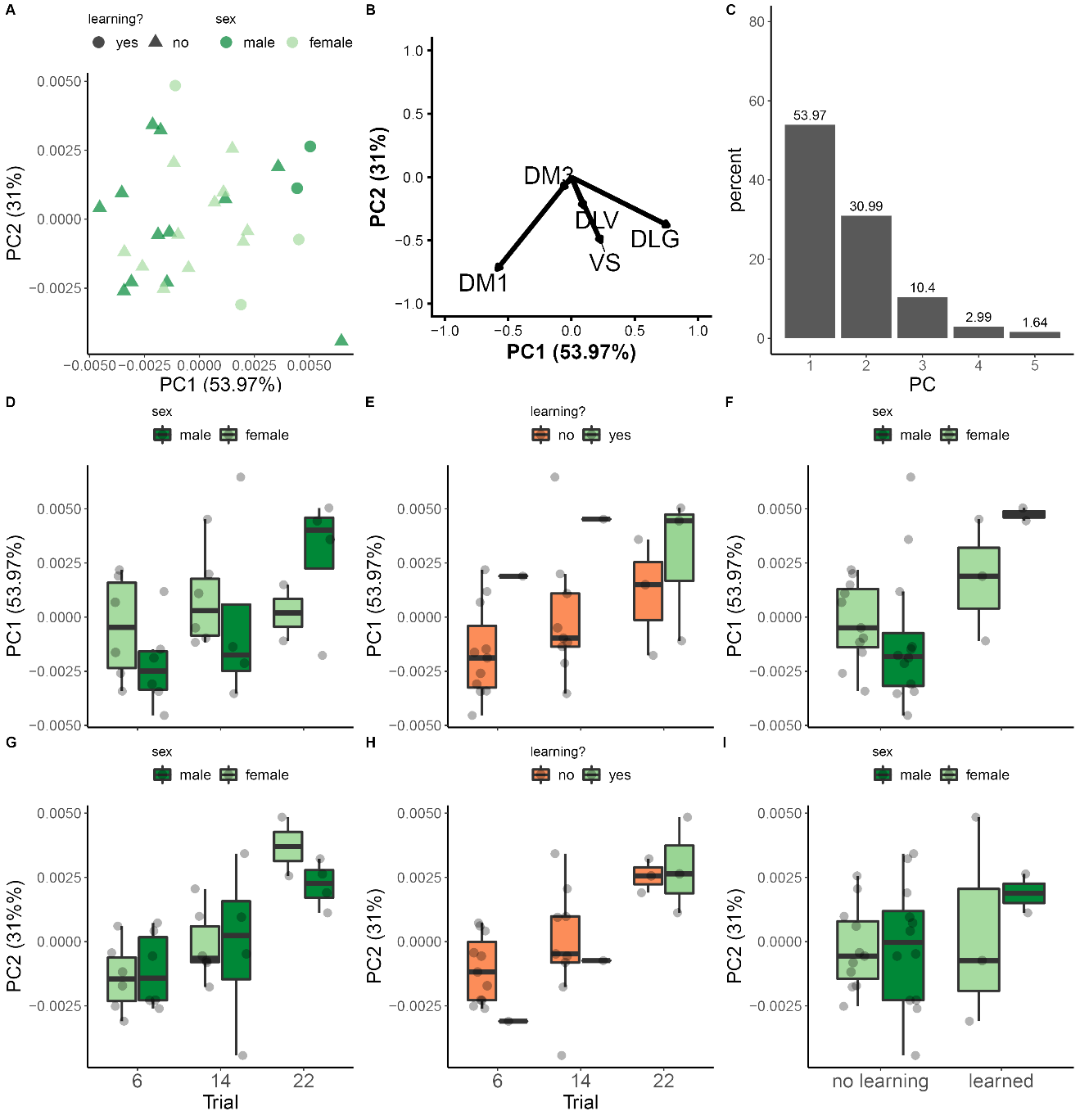

*Supplemental Figure 6*. Principal component analysis (PCA) of neural activity patterns in a non-social context. A) Scatter plot of Fos expression data does not pull out differences by sex on either PC, however PC1 separated out the Dm-1 and Dl-g brain regions (B). C) Plot showing the percent of the variance explained by each PC. PC1 (D-F) and PC2 (G-I) loadings plotted across trials based on the sex of individuals (D and G) and whether learning response was reached (E and H).

**Supplemental Tables**

| % group response | *X^2^* | *p* value |
| --- | --- | --- |
| 50% | 3.5 | 0.173 |
| 60% | 3.5 | 0.175 |
| 70% | 3.5 | 0.175 |
| 80% | 4.3 | 0.118 |
| 100% | 2.8 | 0.252 |

*Supplemental Table 1*. Log-rank test comparisons between percent of group response. There was no statistical difference between log-rank tests in the cumulative response probability based on the percent of the group required to correctly respond at a given trial.

|  |  | Trial of brain collection | | |
| --- | --- | --- | --- | --- |
| SOCIAL CONTEXT | *Observer status* | *Trial 6* | *Trial 14* | *Trial 22* |
|  | Dominant males | n = 4 | n = 3 | n = 4 |
|  | Subordinate males | n = 4 | n = 3 | n = 4 |
|  | Females | n = 4 | n = 3 | n = 4 |
|  | Total per trial | n = 12 | n = 9 | n = 12 |
| NON-SOCIAL CONTEXT | *Individual sex* | *Trial 6* | *Trial 14* | *Trial 22* |
|  | Males | n = 6 | n = 4 | n = 4 |
|  | Females | n = 6 | n = 6 | n = 2 |
|  | Total per trial | n = 12 | n = 10 | n = 6 |

*Supplemental Table 2*. Experimental design and brain collection. Breakdown of number of individuals (based on status) taken immediately following trials 6, 14, and 22 of the association task in both context.

|  | *Factors* | *Df* | *Sums of Squares* | *Mean Square* | *F-value* | *Significance* |
| --- | --- | --- | --- | --- | --- | --- |
| *Dl-g* | trial | 2 | 0.00382 | 0.000191 | 8.878 | *P < 0.001* |
|  | context | 1 | 0.00438 | 0.00438 | 203.754 | *P < 0.001* |
|  | trial*context | 2 | 0.000095 | 0.000048 | 2.22 | *P* = 0.114 |
| *Dl-v* | trial | 2 | 0.0000214 | 0.0000107 | 1.197 | *P* = 0.307 |
|  | context | 1 | 0.0018127 | 0.0018127 | 202.793 | *P < 0.001* |
|  | trial*context | 2 | 0.0000144 | 0.0000072 | 0.807 | *P* = 0.45 |
| *Dm-1* | trial | 2 | 0.0027752 | 0.0013876 | 54.63 | *P < 0.001* |
|  | context | 1 | 0.0030711 | 0.0030711 | 120.92 | *P < 0.001* |
|  | trial*context | 2 | 0.006079 | 0.0003040 | 11.97 | *P < 0.001* |
| *Dm-3* | trial | 2 | 0.0000511 | 0.0000256 | 1.573 | *P =* 0.213 |
|  | context | 1 | 0.0015360 | 0.001536 | 94.479 | *P < 0.001* |
|  | trial*context | 2 | 0.0000328 | 0.0000164 | 1.01 | *P =* 0.368 |
| *Vs* | trial | 2 | 0.00097 | 0.000485 | 8.776 | *P < 0.001* |
|  | context | 1 | 0.005267 | 0.005267 | 95.324 | *P < 0.001* |
|  | trial*context | 2 | 0.000453 | 0.000227 | 4.103 | *P = 0.0197* |

*Supplemental Table 3*. Analysis of variance results for neural activity patterns by context and trial across brain regions. There was a significant main effect of trial in Dl-g, Dm-1, and Vs regions, and a significant main effect of learning context (social v non-social) across all regions. There was an interaction effect between trial and learning context in the Dm-1 and Vs.

|  | *Factors* | *Df* | *Sums of Squares* | *Mean Square* | *F-value* | *Significance* |
| --- | --- | --- | --- | --- | --- | --- |
| *Dl-g* | learning | 1 | 0.0022078 | 0.0022078 | 115.6 | *P < 0.001* |
|  | context | 1 | 0.0027931 | 0.0027931 | 146.3 | *P < 0.001* |
|  | learning*context | 1 | 0.0000363 | 0.0000191 | 1.9 | *P* = 0.171 |
| *Dl-v* | learning | 1 | 0.001757 | 0.0001757 | 17.571 | *P < 0.001* |
|  | context | 1 | 0.0015537 | 0.0015537 | 155.402 | *P < 0.001* |
|  | learning*context | 1 | 0.00000298 | 0.0000028 | 0.276 | *P* = 0.601 |
| *Dm-1* | learning | 1 | 0.000456 | 0.000456 | 10.210 | *P < 0.001* |
|  | context | 1 | 0.003925 | 0.003925 | 87.837 | *P < 0.001* |
|  | learning*context | 1 | 0.000230 | 0.00023 | 5.137 | *P =* 0.0257 |
| *Dm-3* | learning | 1 | 0.0002113 | 0.0002113 | 11.738 | *P < 0.001* |
|  | context | 1 | 0.0012132 | 0.0012132 | 67.403 | *P < 0.001* |
|  | learning*context | 1 | 0.0000010 | 0.0000010 | 0.054 | *P =* 0.816 |
| *Vs* | learning | 1 | 0.002696 | 0.002696 | 46.167 | *P < 0.001* |
|  | context | 1 | 0.003419 | 0.003419 | 58.552 | *P < 0.001* |
|  | learning*context | 1 | 0.000175 | 0.000175 | 2.994 | *P = 0.0869* |

*Supplemental Table 4*. Analysis of variance results for neural activity patterns by context and learning response. There were significant main effects of learning (yes v no) and learning context (social v non-social) across all brain regions. There was a significant interaction between learning and learning context in the Dm-1.

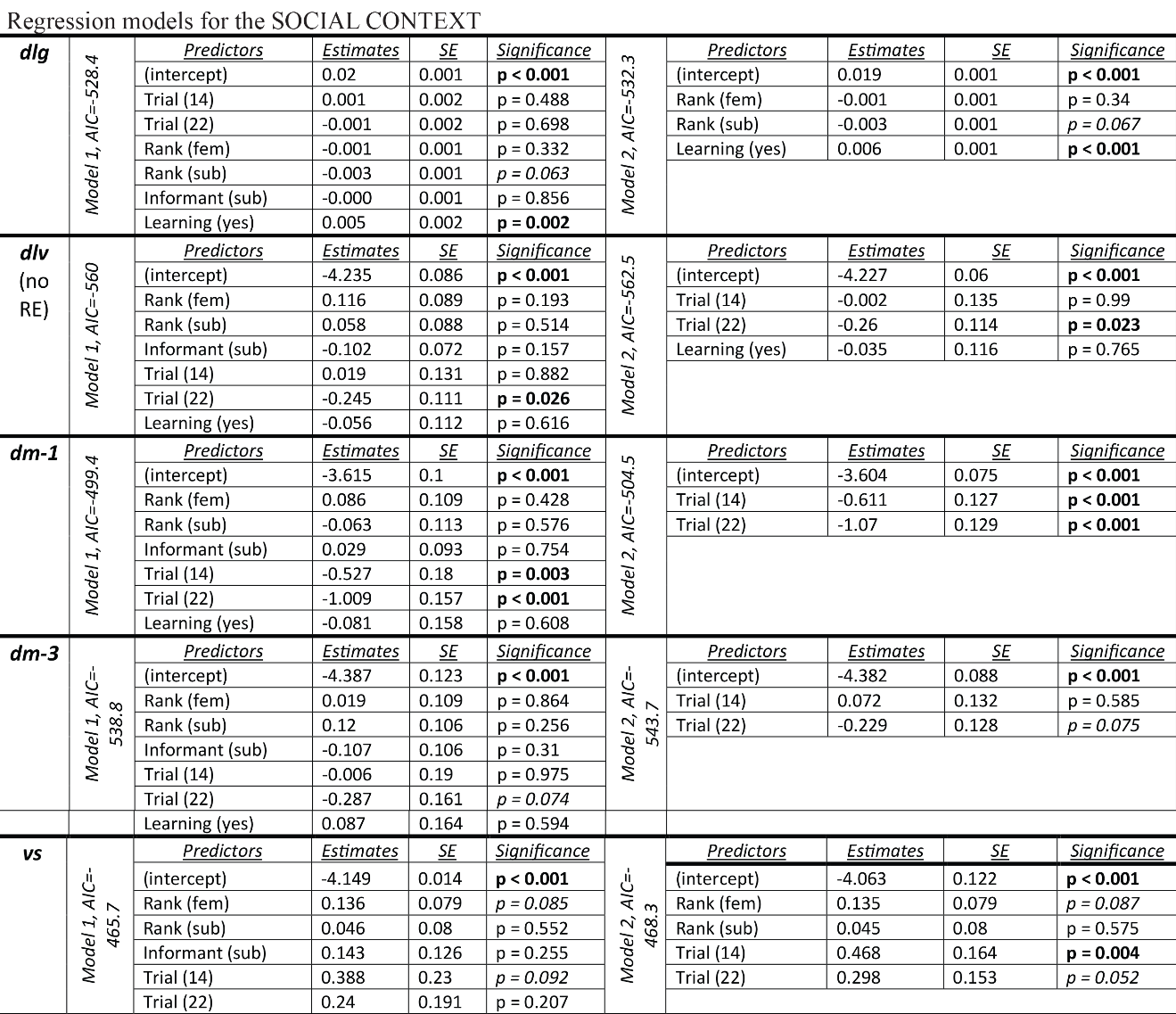

*Supplemental Table 5*. Generalized linear mixed models for each brain region in the social context. We performed iterative model comparisons to examine what factors best predict neural activity in the Dl-g, Dl-v, Dm-1, Dm-3, and Vs separately. In the Dl-g, learning was the best predictor of activity, while trial most significantly predicted activity in the Dl-v. Trial best fit activity in both the Dm-1 and Dm-3, and in the Vs. Across all regions, factors such as individual rank and informant status did not have a significant effect on neural activity.

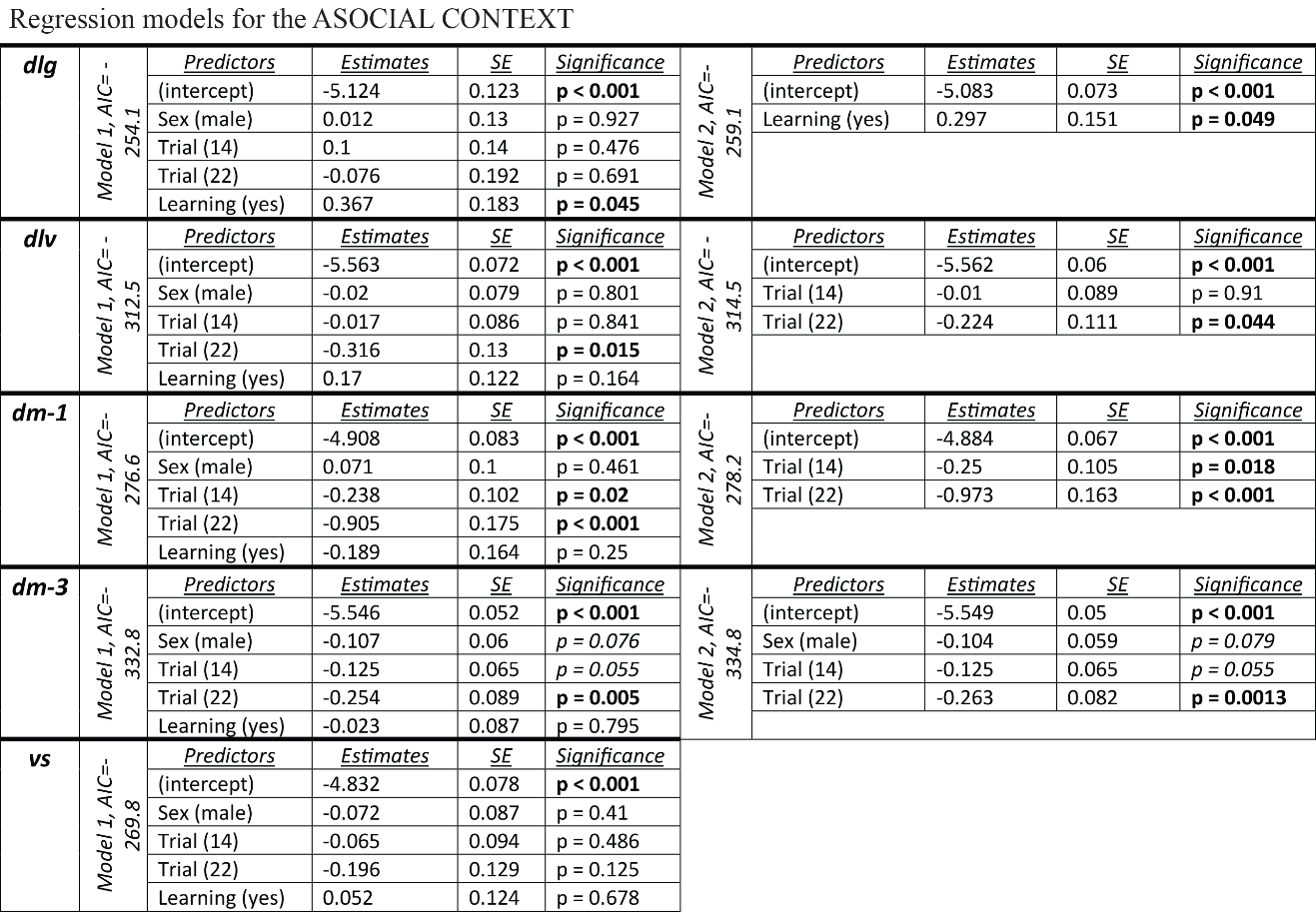

*Supplemental Table 6*. Generalized linear mixed models for the non-social context. We performed iterative model comparisons to examine what factors best predict neural activity in the Dl-g, Dl-v, Dm-1, Dm-3, and Vs separately. In the Dl-g, learning was the best predictor of activity, while trial most significantly predicted activity in the Dl-v. ‘Trial’ best fit activity in both the Dm-1 and Dm-3, and in the Vs. Individual sex did not have a significant effect on neural activity in the non-social context.
